# Genome-wide binding analyses of HOXB1 revealed a novel DNA binding motif associated with gene repression

**DOI:** 10.1101/2020.12.29.424720

**Authors:** Narendra Pratap Singh, Bony De Kumar, Ariel Paulson, Mark Parrish, Carrie Scott, Ying Zhang, Laurence Florens, Robb Krumlauf

**Affiliations:** Stowers Institute for Medical Research, Kansas City, Missouri 64110, USA; University of North Dakota, School of Medicine and Health Sciences, Grand Forks, North Dakota, USA; Department of Anatomy and Cell Biology, Kansas University Medical Center, Kansas City, Kansas 66160, USA

**Author notes:** These authors contributed equally to this work (Narendra Pratap Singh, Bony De Kumar). Corresponding authors: Robb Krumlauf, 1000 E. 50, Bony De Kumar, 1301 N Columbia Road, Stop 9037, Grand Forks, ND 58202-9037.

**Keywords:** Transcription factor, Mouse Hox genes, DNA binding motif, HOXB1, PBX1, REST

## Abstract

Knowledge of the diverse DNA binding specificities of transcription factors is important for understanding their specific regulatory functions in animal development and evolution. We have examined the genome-wide binding properties of the mouse HOXB1 protein in ES cells differentiated into neural fates. Unexpectedly, only a small number of HOXB1 bound regions (7%) correlate with binding of the known HOX cofactors PBX and MEIS. In contrast, 22% of the HOXB1 binding peaks display co-occupancy with the transcriptional repressor REST. Analyses revealed that co-binding of HOXB1 with PBX correlates with active histone marks and high levels of expression, while co-occupancy with REST correlates with repressive histone marks and repression of the target genes. Analysis of HOXB1 bound regions uncovered enrichment of a novel 15 base pair HOXB1 binding motif *HB1RE* <u>(H</u>OX<u>B1</u> <u>r</u>esponse <u>e</u>lement). *In vitro* template binding assays showed that HOXB1, PBX1 and MEIS can bind to this motif. *In vivo,* this motif is sufficient to direct expression of a reporter gene and over-expression of HOXB1 selectively represses this activity. Our analyses suggest that HOXB1 has evolved an association with REST in gene regulation and the novel *HB1RE* motif contributes to HOXB1 function in part through a repressive role in gene expression.

## 1. Introduction

The binding of transcription factors (TFs) at specific regulatory sequences in the genome underlies the control of gene expression and regulation of transcriptional networks essential for programming animal development and organogenesis and for modulating physiological and pathological processes [1,2]. In the evolution of vertebrates, there is evidence for multiple rounds of duplication of genes and entire genomes [3–6], which has expanded the number of members in many TF families. The process of gene duplication and divergence in animal evolution provides a mechanism for the origin of novel gene functions. Paralogous genes in highly conserved TF families may undergo changes resulting in altered functional activities and modified interactions between the TFs with their regulatory regions. This diversification of protein function and the adoption of novel roles has been implicated as a major driver in the evolution of phenotypic complexity, diversity and innovation [7–11]. Changes in *cis-*regulatory sequences over evolutionary time are also a major contributing factor to morphological diversity. Because DNA-binding motifs are in general relatively short sequence motifs (~8-15 bp), small changes in specific binding motifs can have a progressive and significant impact on the efficiency of binding by TFs, resulting in modifications of organismal phenotypes [12,13].

Much less is known about how sequence changes in TF proteins impact their functional activities. Amino acid variations in TF sequences may alter their DNA-binding specificity, leading to broad alterations in patterns of gene expression mediated by changes in their interactions with downstream target genes. This may result in the rewiring of circuits in fundamental transcriptional regulatory networks. Studies of conserved families of TFs have shown that they maintain a surprisingly high degree of similarity in their *in vitro* DNA-binding properties over 600 million years of evolution [14–17]. This has been used as a basis to suggest that changes in TFs, which significantly alter their DNA-binding specificities, are generally considered as a less common mechanism underlying the evolution of phenotypic diversity. However, gene duplication in evolution generates multiple copies of TFs that may serve as substrates for preserving ancestral activities and creating new opportunities for functional divergence [17]. Collectively, this process leads to the emergence of families of paralogous TFs which may contain diversified DNA-binding specificities and new functional activities [16,18]. Thus, identifying and characterizing novel DNA binding motifs of functionally diverged TFs can provide insight into the mechanisms of TF evolution and its contributions to the diversification of regulatory networks in animal evolution.

The HOX family of homeodomain-containing transcription factors are a useful model to investigate the evolution of novel DNA binding specificities. *Hox* genes have conserved roles in the specification of the anteroposterior (AP) body axis in a wide range of animals, from invertebrates to vertebrates, and changes in their expression and function are associated with the diversification of animal body plans [10,19,20]. The regulation, expression, and function of *Hox* genes have been extensively examined [6,15,21–25]. However, how different HOX proteins, with similar *in vitro* DNA binding properties, regulate different developmental programs along the AP axis is poorly understood.

In a recent study, we examined this question by using ChIP-seq (Chromatin immunoprecipitation followed by deep sequencing) approaches to characterize the genome-wide binding properties of HOXA1 and HOXB1 proteins in mouse ES cells differentiated into neural fates [26]. These proteins bind to distinct and largely non-overlapping sets of target genes. Analyses of the binding profiles demonstrated that a large number of the HOXA1 bound regions are co-bound with PBX1 and MEIS [26–28], which are known cofactors for many HOX proteins and other TFs [29–31]. In contrast, only a small subset of HOXB1 bound regions display co-occupancy with PBX and MEIS. Analysis of the underlying sequences in the binding peaks revealed that HOXA1 bound regions are enriched for known consensus motifs for PBX and MEIS, while motifs enriched in HOXB1 bound regions have diversified and represent a different set of DNA motifs. The presence of unknown binding motifs in HOXB1 bound peaks is interesting because cross-species functional analyses have demonstrated that mouse HOXA1 has retained the ancestral functions of *Drosophila* Labial, while HOXB1 has diverged and evolved new functions [26]. Therefore, investigating the properties of novel HOXB1 binding motifs can provide insight into the properties associated with its neofunctionalization.

In this study, we have examined enriched motifs associated with genome-wide binding profiles of the mouse HOXB1 protein and investigated their properties and regulatory potential. We focused on the characterization of one of the novel motifs enriched in the HOXB1 bound regions and named this 15-base pair motif *HB1RE* <u>(H</u>OX<u>B1</u> <u>r</u>esponse <u>e</u>lement). The center of this motif has a known *in vitro* HOX binding sequence (5’-ATTA-3’) [32]. However, the flanking sequences are much longer and are distinct from the previously characterized Hox-PBX bipartite motif [33,34]. This is widely distributed across the genome. Using an *in vitro* template binding assay and a multidimensional protein identification technology (MudPit)-based mass-spectrometry approach, we found that PBX1 can bind to the *HB1RE* motif in nuclear extracts. Genomic analyses revealed that different subsets of genomic regions enriched for this *HB1RE* motif are also bound by PBX1 or REST proteins. Functional analysis of this motif using transgenic constructs in electroporated chicken embryos showed that multiple copies of this motif are sufficient to drive expression of a reporter gene in the neural tube, suggesting it has *in vivo* regulatory potential. Over-expression of HOXB1 in this transgenic assay selectively represses reporter activity mediated by this motif. Together our analyses suggest that HOXB1 binds the *HB1RE* motif and modulates *in vivo* function in part through a repressive role in gene expression.

## 2. Materials and Methods

### 2.1. ES cell culture and induction of KH2 cells with retinoic acid

KH2 mouse ES cells [35] at Passage 12 were cultured on gamma-irradiated feeder cells with DMEM containing 15% fetal bovine serum, NAA and β-mercaptoethanol. The media was changed 3h before passaging. In transfer to feeder-free conditions, the media was aspirated and washed twice with PBS, then 2 ml of pre-warmed Trypsin/EDTA solution was added and placed in an incubator at 37°C for 1 m. During this period colonies float off when tapping the plate. Trypsin activity was stopped by adding 5 ml of FCS-ES medium to the flask. Colonies were dissociated into single cells by pipetting up and down several times and pelleting the cells by centrifugation at 1000 rpm for 5 m. Media was aspirated and the cells were resuspended in an appropriate volume of fresh ES cell medium. Gelatinized plates were used for the feeder-free culture of the ES cells. Culture plates were treated 30 m before seeding with 0.1% Gelatin. Gelatin was later aspirated just before seeding of the cells. In feeder-free culture, the KH2 cell lines were grown using N2B27+2i media supplemented with 2,000 U/mL of ESGRO (Millipore) [36]. Cells were differentiated on a gelatinized plate using differentiation media (DMEM + 10% (vol/vol) Serum + NEAA + 3.3 μM RA). ES cells were harvested at 80–90% confluency for the experiments. In genomic experiments, mouse ES cells were differentiated using retinoic acid (RA) for 24 h [37]. In the case of the HOXB1 epitope tagged KH2 line, doxycycline was added with RA to simultaneously differentiate the cells and induce expression of HOXB1.

### 2.2. ChIP-seq and ATAC-seq

Anti-Flag antibody was used to pull down 3XFLAG tagged-HOXB1 for ChIP-seq experiments. We used the Upstate protocol for ChIP-seq experiments as described in Smith *et al* [38]. All other ChIP-seq experiments were done using specific antibodies against the transcription factors (Anti-PBX: Santacruz; SC-888, Anti-MEIS: Santacruz; SC-25412, Anti-REST: Abcam; ab21635) and histone proteins (Anti-H3K27Ac: Abcam; ab4729, Anti-H3K4me3: Abcam; ab8895). Expression analysis of the genes at various time points (0, 4, 6, 12, 24 and 36 hour) after RA differentiation of ES cells were analysed using RNA-seq [39]. Buenrostro and colleagues protocol was used for the chromatin accessibility (ATAC-seq) experiment using approximately 50,000 feeder-free uninduced and differentiated ES cells [40]. The raw data files for ChIP-seq, ATAC-seq, RNA-seq data have been submitted to the NCBI BioProject database: https://www.ncbi.nlm.nih.gov/sra/?term=PRJNA503882, <u>PRJNA341679</u>, PRJNA335616, <u>GSE61590</u>, Sequence Read Archive under accession no: SRX4980243 to SRX4980246 and Gene Expression Omnibus (GEO; http://www.ncbi.nlm.nih.gov/geo/) under series accession number GSE61590. The original data is also deposited to Stowers Institute Original Data Repository and available online at http://odr.stowers.org/websimr/.

### 2.3. Genomic analysis

Sequencing reads were aligned to the mouse genome (mm10) with bowtie2 using default parameters [41]. Peaks were called with MACS2 [42] with parameters “-p 0.25 -m 5 50” to ensure sufficient peaks for IDR analysis. MACS2 peak coordinates were trimmed back to meet the actual IP signal. To compare the peaks from replicates we used irreproducibility discovery rate (IDR version 2.0.7) and analyzed top 100,000 peaks by p-value, and only paired peaks with MACS2 fold-change >= 4, q-value <= 0.05, and IDR global p <= 0.1 were collected for future analyses.

For analysis of gene expression levels for near adjacent genes in HOXB1 binding regions we used our previously published RNA-seq data detailing a time-course of changes in transcriptional profiles during differentiation of ES cells [39]. In brief, RNA was isolated using TRIzol reagent. RNA sample with more than 9 RIN (RNA-integrity number) was used for library preparation. PolyA-selected RNA libraries were prepared using mRNA-seq Library Prep Kit (Illumina, RS-100-0801) according to the manufacturer’s protocol (15018460 Rev AOct 10). 1 μg Total RNA was enriched for poly(A)+ RNA by oligo(dT) selection. The purified libraries were quantified with the High Sensitivity DNA assay on an Agilent 2100 Bioanalyzer and. Libraries were sequenced single read with 36 nt sequencings on a GAIIx, and fastq files were returned. For each sample, reads were aligned to mm10 using Hisat2, allowing uniquely aligning reads only. Differential Gene expression analysis was done using DeSeq2. Different time points were compared against the gene expression profile of Uninduced ES cells.

DNA motif for each HOXB1 peak list with kmer contents (k) from 4-12 was examined, and any peak where any single kmer occupied >= 30% of width was excluded. The remaining top 1000 peaks by p-value, widths within 200-500bp, were selected, and summit ±50bp windows were sent to MEME 4.12.0, parameters “-nmotifs 50 -minw 6 -maxw 25 -mod zoops -evt 10 -maxiter 200 -minsites 50 -maxsites 1000 -maxsize 1E9” [43]. Resulting motifs were compared to Transfac release 2019.3 [44] and JASPAR 2018 [45] vertebrate motifs with Tomtom 4.12.0, parameters “-min-overlap 4 -thresh 0.1”. Motifs were scanned with FIMO 4.12.0, parameters “--bgfile --motif-- --thresh 0.0001”, and further filtered for q-value <= 0.1. Motif search spaces were the summit ±50bp windows for the entire peaklist, and 10 shuffled copies as the background. Enrichments were calculated with Fisher’s Exact test, BH-adjusted p-values <= 0.05. All HOXB1 peaks matching any of the three HRE variants (original, dreme-full, dreme-core) using FIMO at p<=1e-4 were selected (188 peaks) [46].

Heatmaps were generated using CoverageView package from Bioconductor using 5kb windows centered on peak summit, averaged into 100 bins. Log fold change values between IP and input samples were calculated using reads per million (RPM) and only positive values are shown. For the HOXB1 peak heatmap, the data shown is the highest-LFC data from any constituent HOXB1, thus the heatmap column shows the “best-case scenario” for the HOXB1 peak list. The nearest protein-coding neighbors were identified using Ensembl 91 and then KEGG (Kyoto Encyclopedia of Genes and Genomes) pathway analyses was performed on these sets of genes [47].

### 2.4. Template binding assay

This assay was done using a protocol for nucleosome binding assay on a template with few modifications mentioned below [48,49]. 5′ –biotinylated 6X *HB1RE* motif DNA fragments were bound to streptavidin dyna beads (75 fmol DNA/μL beads) in 200 μL of binding buffer (20 mM Hepes, PH 7.9, 0.05% NP-40, 10% Glycerol, 10 mM MgCl2, 2 mM DTT, 0.5 mM PMSF, 100 μg/ml BSA, ddh20. In parallel, 10 μl of nuclear extract from 24h differentiated 3xFLAG-HOXB1 transgenic ES cells was incubated with 90 μL of reaction buffer (10 mM HEPES (pH7.9), 0.1 M NaCl, 1.5 mM MgCl_2_, 0.05% TritonX-100 w/protease inhibitor cocktail + 20 μl re-constitution buffer (1 mM ATP, 50 μM ZnCl2, 45 ng/μl sheared lambda phase DNA) and NaCl (varies as per reaction requirement) for 10 m in 30°C vibrating incubator at 1200 rpm to remove non-specific binding. Then 5 μl of DNA-bead complex was added to the reaction and incubated for 20 m in a 30°C vibrating incubator at 1200 rpm. Template bound intermediates were separated from supernatant fractions by magnetic racks. Following fractionation, beads were washed with 200 μl of reaction buffer, transferred to a fresh microcentrifuge tube, and bound proteins were eluted with 1x SDS sample buffer. Template bound intermediates were analyzed by western blot using infrared antibody imaging (Li-COR Biosciences) or MudPIT.

### 2.5. Western Blot

A standard western blotting protocol was used to detect HOXB1 in template binding assay. Anti-FLAG antibody (Sigma F-1804) was used to detect the 3xFLAG tagged HOXB1 protein in the eluted samples.

### 2.6. Multidimensional Protein Identification Technology (MudPIT)

TCA-precipitated proteins were urea-denatured, reduced, alkylated and digested with endoproteinase Lys-C (Roche) followed by modified trypsin (Promega) [50,51] Peptide mixtures were loaded onto 250 μm fused silica microcapillary columns packed with strong cation exchange resin (Luna, Phenomenex) and 5-μm C18 reverse-phase (Aqua, Phenomenex), and then connected to a 100 μm fused silica microcapillary column packed with 5-μm C18 reverse-phase (Aqua, Phenomenex) (Florens and Washburn, 2006). Loaded microcapillary columns were placed in-line with a Quaternary Agilent 1100 series HPLC pump and an LTQ linear ion trap mass spectrometer equipped with a nano-LC electrospray ionization source (ThermoScientific, San Jose, CA). Fully automated 10-step MudPIT runs were carried out on the electrosprayed peptides, as described in [50]. Tandem mass (MS/MS) spectra were interpreted using ProluCID (Xu et al., 2015) against a database consisting of 61441 non-redundant *M. musculus* proteins (NCBI, 2020-11-03 release), 426 usual contaminants (human keratins, IgGs, and proteolytic enzymes). To estimate false discovery rates (FDR)s, the amino acid sequence of each non-redundant protein entry was randomized, resulting in a total search space of 123728 non-redundant sequences. All cysteines were considered as fully carboxamidomethylated (+57 Da statically added), while methionine oxidation was searched as a differential modification. DTASelect [52] and swallow, an in-house developed software, were used to filter ProLuCID search results at given FDRs at the spectrum, peptide, and protein levels. Here all controlled FDRs were less than 1%. All 5 data sets were contrasted against their merged data set, respectively, using Contrast v 1.9 and in house developed sandmartin v0.0.1. Our in-house developed software, NSAF7 v0.0.1, was used to generate spectral count-based label-free quantitation results. The MS datasets may be obtained from the MassIVE repository with accessions MSV000086596 (with password: NPS01146) and from the ProteomeXChange with accessions PXD023129.

### 2.7. Reporter assays in electroporated chicken embryos

*In ovo* electroporations of chicken embryos were performed as previously described [53–55]. The *Hoxb1* auto-regulatory element (*ARE)* [56] and *HB1RE* were cloned into the β-galactosidase reporter vector *BGZ40* (1μg/μl) [57] and was injected into the neural tube of 7–10 somite(s) or 22s–27s stage chick embryos either alone or in combination with DNA containing (1μg/μl) of specific *Hox* cDNAs cloned into the pCMS expression vector. Injected embryos were then electroporated (3 pulses of 18 V, 50 ms On, 950 ms off) and allowed to develop for an additional 18-24 h. After harvesting, embryos were washed in PBS 0.02% NP40 and fixed for 1h in 1% paraformaldehyde, 0.2% glutaraldehyde, 2 mM MgCl2, 5 mM EGTA and 0.02% NP40 in PBS at 4°C. After fixation, the embryos were washed 3 x 20 m with rocking in PBS+0.02% NP40. Embryos were then stained in 5 mM K3Fe(CN)6, 5 mM K4Fe(CN)6-3H20, 2 mM MgCl2, 0.01% Sodium Deoxycholate, and 0.02% NP40 + 1mg/ml Xgal at 4°C in the dark for 12-36 h. Staining was observed and captured on a Leica MZ APO stereoscope/camera/computer setup.

## 3. Results

### 3.1. Characterizing the genome-wide binding patterns of HOXB1 in mouse ES cells

Generating specific antibodies against specific HOX proteins is challenging because of the presence of highly conserved regions in the proteins. To overcome this limitation, we generated an epitope-tagged (3XFLAG) of HOXB1 in the mouse KH2 ES cell line by inserting the coding region fused with the epitope at a defined endogenous locus designed for tight doxycycline inducible expression [27,35,36,39]. These modified KH2 ES cells were then differentiated into neuroectodermal cell fates using previously established retinoid treatments, along with doxycycline to induce expression of the epitope-tagged version of HOXB1 [37,39]. The expression of HOXB1 from inducible locus was conducted under conditions that optimized the amount of doxycycline to generate levels comparable to the endogenous *HOXB1* gene [27,39]. Gene expression analyses of the differentiated ES cells over a time-course revealed that the transcription profile at 24 h of differentiation is similar to that of the mouse embryonic hindbrain and spinal cord. Therefore, we used cells differentiated cells for 24 h to map the genome-wide binding targets of HOXB1 using Chromatin immunoprecipitation followed by deep sequencing (ChIP-seq) with an antibody against the epitope tag [26,27].

Using replicate samples in a series of ChIP-seq experiments we reproducibly mapped 2058 binding peaks for HOXB1 in the mouse genome (Figure 1A) [26]. Gene ontology (GO) and KEGG (Kyoto Encyclopedia of Genes and Genomes) pathway analysis of nearest neighbor genes in HOXB1 bound peaks suggest that HOXB1 downstream targets are enriched for genes and processes required for neurogenesis, neuronal processes, and behavior (Figure 2A). This is consistent with functional and genetic studies in zebrafish, mouse and humans which demonstrated key functional roles for *HOXB1* in the patterning the hindbrain and facial nerve (VIIth cranial nerve) during craniofacial development [58–61].

**Figure 1:**
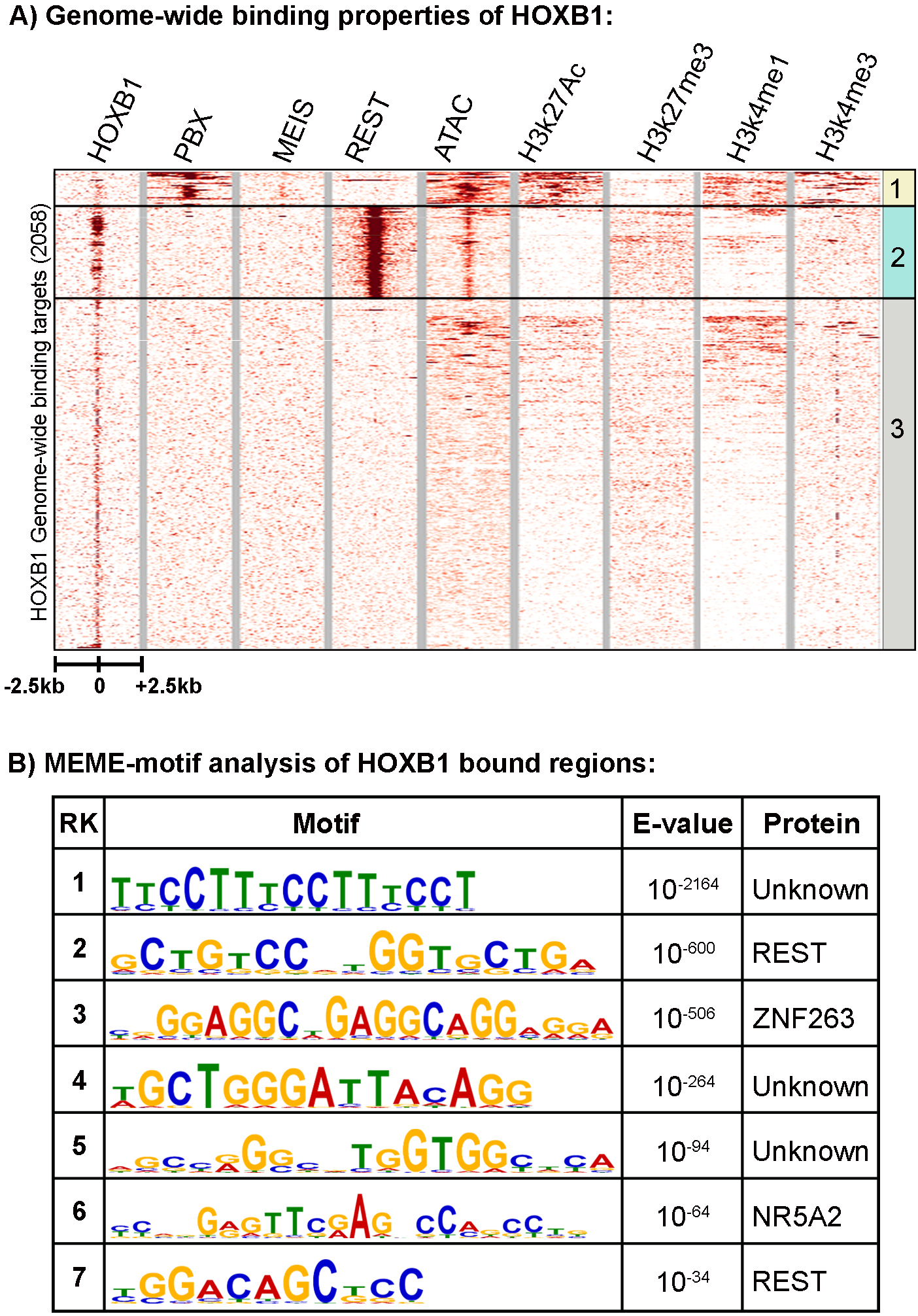
Genome-wide binding pattern of HOXB1: A) Heatmap shows all HOXB1 bound peaks in the genome with ± 2.5Kb flanking region and its overlap with PBX1, MEIS, REST, ATAC-seq (open chromatin), H2k27Ac, H3k27me3, H3k4me1 and H3k4me3. The HOXB1 peaks enriched with PBX1 (top) are boxed as group 1, HOXB1 peaks enriched with REST (middle) are boxed as group 2 while all other peaks are boxed as group 3. B) Top seven motifs identified by MEME (Multiple Expression motifs for Motif Elicitation), motif discovery tool are ranked on the basis of enrichment value (E value). The PWM (position wait matrix) motif logo and name of the known transcription factors are mentioned in the table.

**Figure 2:**
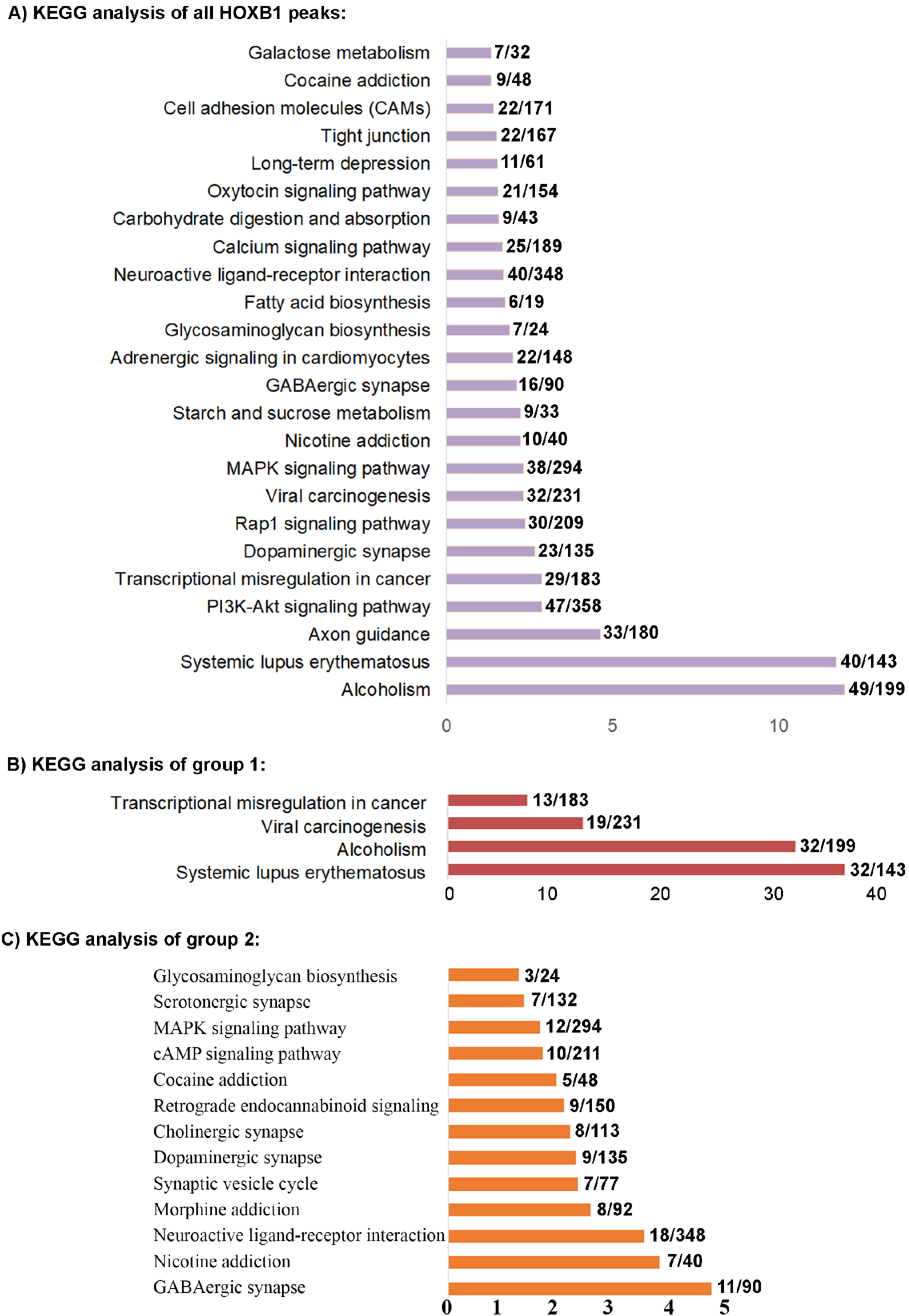
KEGG analysis of HOXB1 peaks: A) KEGG analysis of nearest neighbor genes of all the HOXB1 peaks. B) KEGG analysis of nearest neighbor genes of the HOXB1 peaks overlapping with PBX1 (group 1). C) KEGG analysis of nearest neighbor genes of the HOXB1 peaks overlapping with REST (group 2).

*Cis*-regulatory studies have demonstrated that HOXB1 partners with the HOX co-factors PBX and MEIS, to auto-regulate its expression and cross-regulate several other *Hox* genes during hindbrain segmentation [56,57,62–65]. Hence, we directly compared the genome-wide binding patterns of HOXB1 with that of PBX1 and MEIS and found that only a small number of the total HOXB1 binding peaks (152/2058: group 1) are co-bound with PBX1 and/or MEIS (Figure 1A). This pattern is distinctly different from its paralog HOXA1, which displays significant overlaps in co-occupancy with PBX and MEIS [27], and previous observations on the interactions of PBX and MEIS with other HOX proteins in modulating their DNA binding specificity and function [33,66]. This difference in binding patterns is consistent with the idea that the HOXB1 protein has diverged from HOXA1 and Labial in part through altered interactions with PBX proteins [26].

Analysis for the enrichment of putative transcription factor binding motifs in the regions occupied by HOXB1 uncovered motifs known to bind REST, ZNF263 and NR5A2 proteins, along with several novel/unknown motifs (Figure 1B and ref [26]). In agreement with the lack of co-occupancy of PBX and MEIS the in ChIP-seq data, we find no evidence for enrichment of PBX and MEIS binding motifs. The significant enrichment of consensus motifs for the REST protein complex is interesting, as REST functions as a transcriptional repressor to restrict the expression of neuronal genes in non-neural cells [67–69]. To examine whether REST displays occupancy on these putative motifs, we performed ChIP-seq experiments with antibodies against the REST protein in differentiated ES cells. The heatmap in Figure 1A shows that 22% (462/2058, group 2) of HOXB1 peaks are co-bound by the REST transcriptional repressor complex, validating the motif analyses. This raises the possibility of functional interactions between HOXB1 and the REST co-repressor complex in the regulation of downstream target genes.

Most of the HOXB1 bound regions that display co-occupancy with PBX1 (group 1) do not overlap with those co-bound by REST (group 2) and vice versa, suggesting a mutually exclusive relationship between these classes of binding regions (Figure 1A). The remaining peaks bound by HOXB1 form group 3 (1444/2058) and they represent the majority (~70%) of bound regions. Peaks in group 3 do not correlate with binding by PBX, MEIS or REST, implying that they have very different features.

### 3.2. Analysis of the HOXB1 binding peaks and co-association with PBX and REST

Assays for histone marks indicative of epigenetic states and chromatin accessibility (ATAC-seq) were conducted to investigate the features of chromatin-associated with occupancy of HOXB1 and co-binding with PBX1 and REST (Figure 1A). HOXB1 peaks that correlate with PBX1 binding (group 1) display open chromatin in ATAC-seq analysis and enrichment of H3K27Ac, H3K4me1 and H3K4me3 histone marks, known to be associated with active loci. While HOXB1 and REST co-bound regions (group 2) have open chromatin, they lack histone marks indicative of active loci and display a modest enrichment for the H3K27me3 histone mark, known to be associated with Polycomb mediated repression. This suggests that HOXB1 co-occupancy with PBX1 is correlated active regions of the genome, while co-binding with REST is correlated with repressed regions of the genome. Interestingly, the regions that do not display co-occupancy with either PBX1 and REST proteins (group 3) generally have a closed chromatin state and most of the regions are devoid of any histone marks. Although a few of the peaks in cluster 3 display H3K4me1 marks and also have open chromatin, suggesting that they may be poised for activity (Figure 1A).

We further investigated the gene regulatory potential of PBX and REST protein co-binding with HOXB1. We analyzed the expression of differentiating ES cell at various time points and correlated their expression with HOXB1 co-binding with PBX1 and REST (Figure 3 and 4). We found that HOXB1 and PBX1 co-binding at promoter and gene body is associated with higher expression of the genes as compared to genes bound only with HOXB1. In contrast, HOXB1 co-bound regions with REST at promoters and gene bodies show lower expression of the genes as compared to the genes bound only with HOXB1. This suggests that PBX1 binding on HOXB1 bound regions may lead to activation of the locus for gene expression while REST binding suppresses this activity. This also correlates with the known regulatory activities of the PBX1 in gene activation and REST in gene suppression and suggest that HOXB1 may use these proteins as a cofactor for the regulation of target genes.

**Figure 3:**
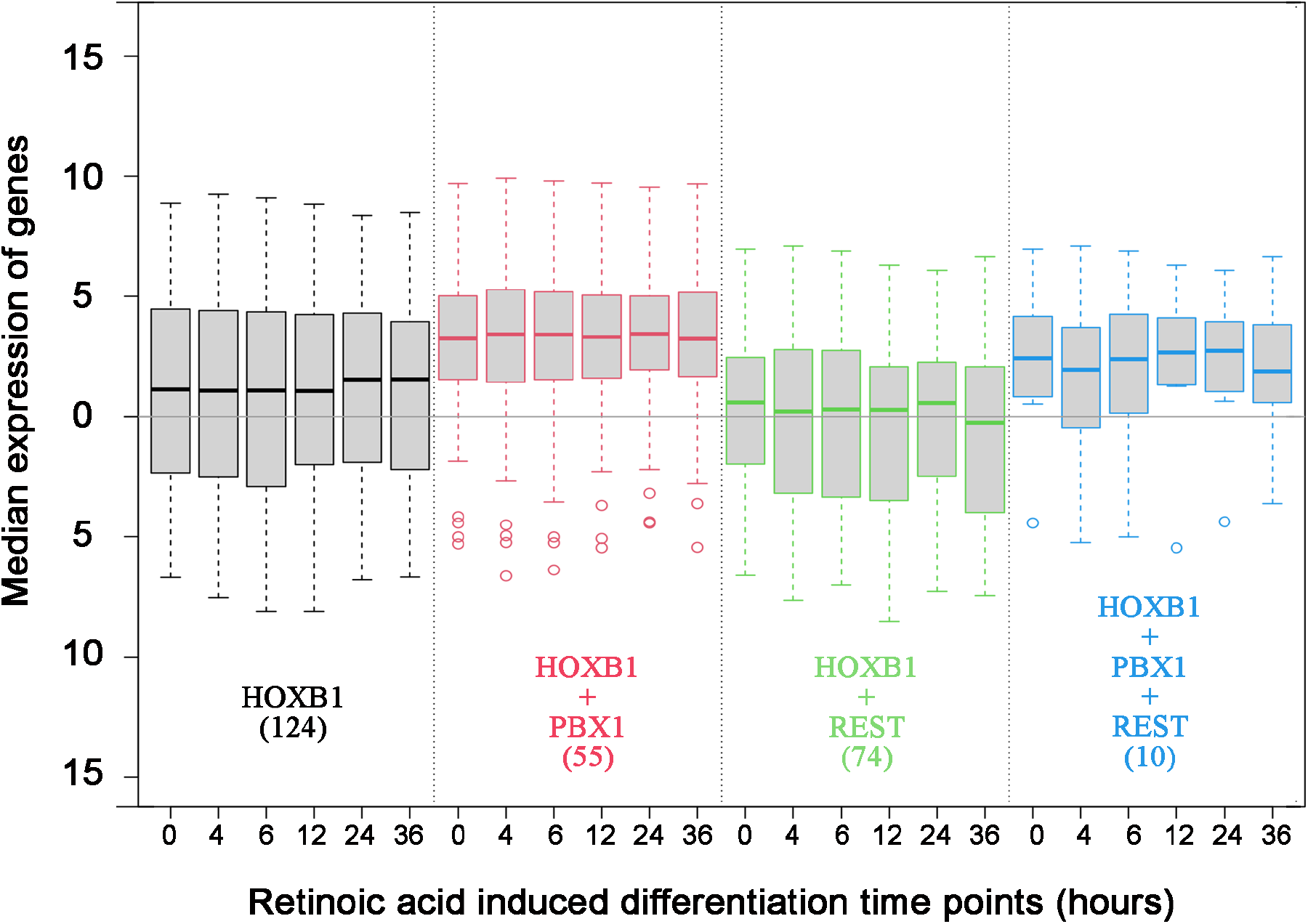
Expression of genes having HOXB1, PBX1 and REST binding at promoter: Box plot shows median expression level of genes (Y axis) in differentiating ES cells at six time points mentioned in hours (X axis). The analysis is done with the four sets of genes separated on the basis of HOXB1 (black), HOXB1 + PBX1 (red), HOXB1 + REST (green) and HOXB1+PBX1+REST (blue) binding at the promoter region. The total number of genes in each set are mentioned in the brackets.

**Figure 4:**
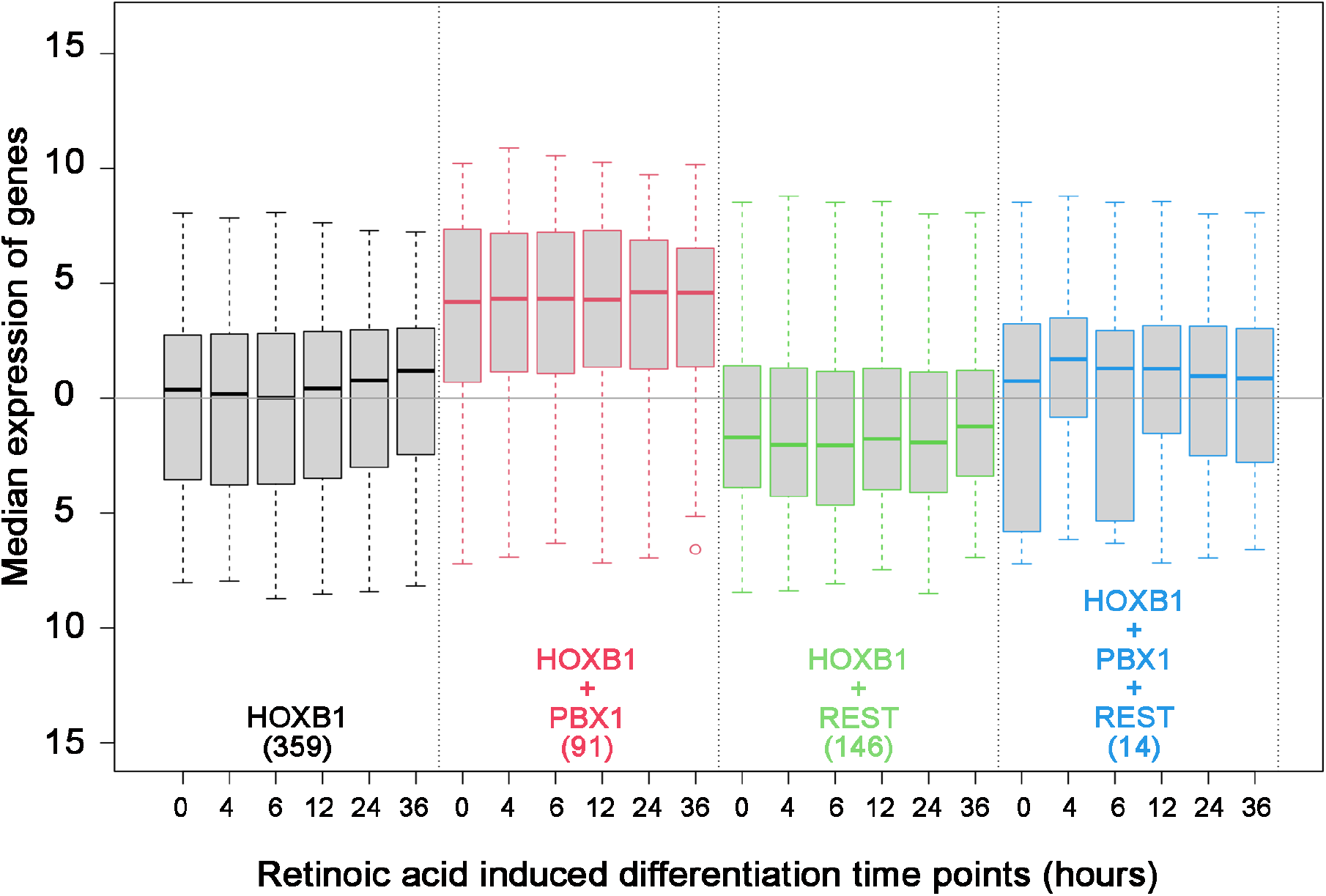
Expression of genes having HOXB1, PBX1 and REST binding at gene body: Box plot shows median expression level of genes (Y axis) in differentiating ES cells at six time points mentioned in hours (X axis). Analysis is done on the four sets of genes separated on the basis of HOXB1 (black), HOXB1 + PBX1 (red), HOXB1 + REST (green) and HOXB1 + PBX1 + REST (blue) binding at the gene body. The total number of genes in each set are mentioned in the brackets.

To explore the potential the function of genes regulated by HOXB1 in groups 1 and 2 associated with co-occupancy of PBX1 or REST, we extended KEGG analysis of the neighboring genes to these groups (Figure 2B and 2C). While the KEGG analysis of all HOXB1 peaks shows enrichment for neurogenesis, neuronal processes, and behavior there is some representation for genes associated with metabolism, biosynthesis, and cancer (Figure 2A). In contrast, the group 1 peaks, where HOXB1 and PBX1 display co-occupancy, associate with genes required in cancer and alcoholism, while the group 2 regions with co-binding of HOXB1 and REST, associate with genes required in neurogenesis, neuronal processes, and behavior (Figure 2B and 2C). This suggests that HOXB1 binding may be associated with different co-factors and co-regulators to modulate the expression of the different sets of genes involved in these distinct processes. Analysis of the group 3 HOXB1 binding peaks, which that do not overlap with either PBX1 or REST occupancy, indicates they appear to lie in gene deserts and are associated with only a small subset of genes involved in axon guidance.

### 3.3. Characterization of an unknown HOXB1 binding target motif, *HB1RE*

To begin to explore properties associated with the neofunctionalization and diversification of HOXB1, we further examined the novel DNA motifs enriched in the HOXB1 bound regions (Figure 1B). In the top 7 enriched motifs, we found a high level of enrichment for three unknown motifs. The top enriched motif is a low complexity motif of 5’-TTTCC-3’ nucleotide repeat. The second unknown motif, ranked 4^th^ overall, is 15 a nucleotide sequence (5’-T/AGCTGGGATTACAGG-3’). This has a 5’-ATTA-3’ sequence positioned in the middle of the motif, which is a well-characterized common sequence associated with *in vitro* analyses of HOX protein binding [32]. Hence, we hypothesized that this motif might be an interesting consensus sequence for exploring HOXB1 protein binding *in vivo*. We named this motif <u>H</u>OX<u>B1</u> <u>R</u>esponse <u>E</u>lement (*HB1RE*) and it is detected in 188 of the 2058 HOXB1 binding peaks (Figure 5A). To further characterize the *HB1RE* motif we used genomic, biochemical, and functional approaches to examine its potential role in HOXB1 mediated gene regulation.

**Figure 5:**
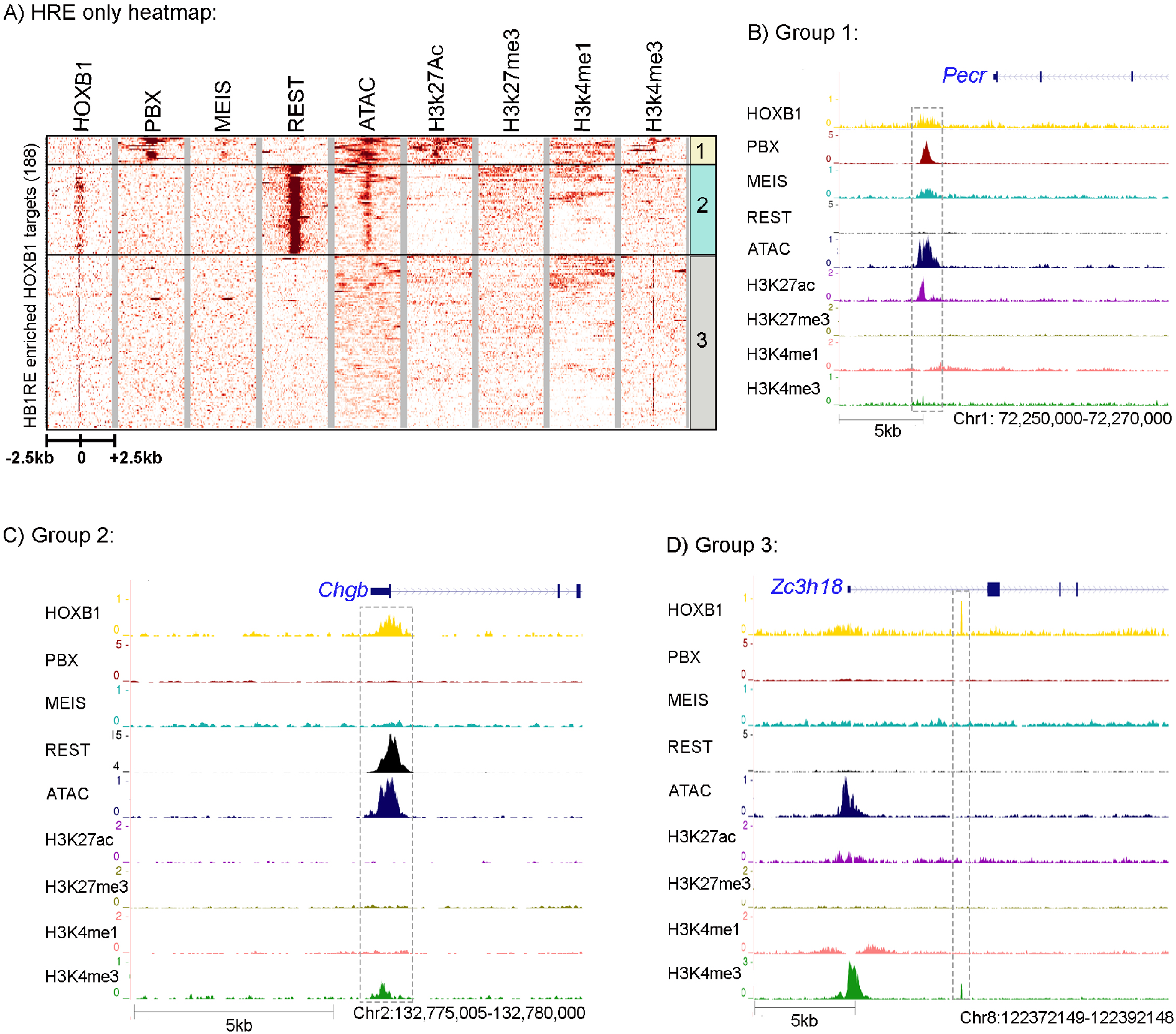
Analysis of HB1RE motif: **A)** Heatmap shows HOXB1 peaks ± 2.5Kb flanking region that is enriched with HB1RE motifs. These peaks are overlapped with PBX1, MEIS, REST, ATAC-seq (open chromatin), H2k27Ac, H3k27me3, H3k4me1 and H3k4me3. The HOXB1 peaks enriched with PBX1 (top) are boxed as group 1, HOXB1 peaks enriched with REST (middle) are boxed as region 2 while all other peaks are boxed as region 3. Genome browser shot of a representative locus for group 1, 2 and 3 are shown in B, C and D sections, respectively.

HOXB1 peaks having the *HB1RE* motif were extracted from the total population of HOXB1 bound regions and compared with the PBX and REST occupancy, open chromatin, and histone marks (Figure 5A). Interestingly, we observed the distribution of this motif in all 3 groups of HOXB1 bound peaks. Of the 188 regions containing the HB1RE motif, 18 falls in group 1 associated with co-occupancy by PBX1 and MEIS while 60 reside in group 2, correlated with co-binding by REST. It is interesting that the group 1 peaks are enriched for co-occupancy of PBX and MEIS, despite not possessing the typical consensus bipartite Hox-PBX motifs associated mediating their physical interactions with HOX proteins and role in modulating DNA binding properties [30,31]. These *HB1RE* containing peaks in group 1 are also enriched for open chromatin and active histone marks, suggesting that HOXB1 and PBX1 may work together in positively regulating gene expression of the adjacent target genes. A browser shot of the *Pecor* locus is presented as an example of loci in this group (Figure 5B). In contrast, the HOXB1 *HB1RE* containing peaks that are co-bound with REST proteins generally do not associate with active histone marks, but they do display open chromatin states. The *Chgb* locus is shown as an example of binding properties in this group 2 class of peaks (Figure 5C). *HB1RE* containing peaks in group 3 show closed chromatin and absence of histone modifications (Figure 5D).

### 3.4. Genomic and biochemical characterization of HB1RE motif

The genomic analyses above imply an association or co-occupancy of HOXB1, PBX1 and REST with the *HB1RE* motif. It is possible that the co-occupancy is direct or indirect. Hence, we used an unbiased approach, based on an *in vitro* template binding assay to identify proteins capable of binding to this motif *in vitro* [48,49]. The template binding assay utilized oligomerized versions of the *HB1RE* motif in combination with nuclear extracts from 24 h differentiated ES cell expressing 3X-FLAG-HOXB1 followed by western blotting or MudPIT (Multidimensional Protein Identification Technology) analysis to identify proteins interacting with the *HB1RE* motif (Figure 6). A western blot using an anti-Flag antibody confirms that HOXB1 binds to this motif (Figure 6B). The MudPIT analysis reveals that PBX1, MEIS1, 2 and 3, are among the most highly enriched interaction proteins compared with control samples (Figure 5C). Since the HB1RE lacks canonical sequences for PBX and MEIS binding, these proteins are likely to bind to non-consensus sequences or are indirectly recruited via interactions with other factors, perhaps HOXB1 itself. These observations along with the genomic analysis indicate that the *HB1RE* motif is capable of interacting with HOXB1, PBX1 and MEIS proteins *in vitro* and *in vivo* to modulate the expression of neighboring genes and represents a new type of Hox target motif. The MudPIT data indicate that several Polycomb complex proteins are also enriched for binding on the *HB1RE* motif, but we do not detect interactions with components of the REST complex. This suggests that the co-occupancy between HOXB1 and REST in peaks containing *HB1RE* motifs is likely to be mediated via the close associate of independent REST binding sites.

**Figure 6:**
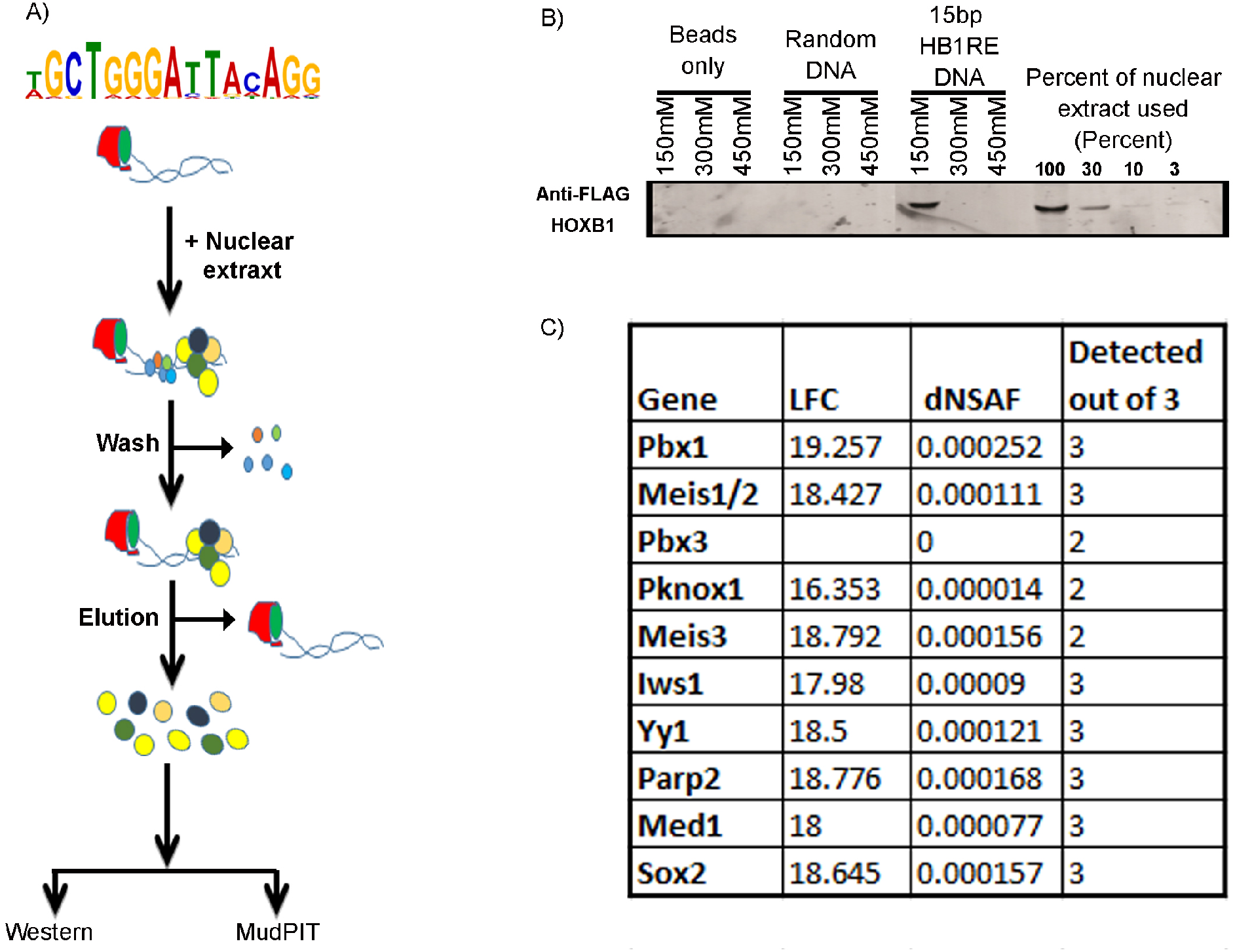
Template binding assay: A) The pictorial representation shows a flow diagram of the template binding assay. B) Western blotting of the eluted samples from template binding assay shows enrichment of HOXB1 only in the HB1RE template and positive control input samples. C) Few known cofactors of HOX proteins identified in MudPIT analysis of the eluted sample is listed with log fold change (LFC), dNSAF (distributed normalized spectral abundance factor) value and detection out of the three replicates.

### 3.5. Reporter assay for regulatory activity of the *HB1RE* motif in chicken embryos

To further investigate the functional properties of the enriched *HB1RE* motif we employed a transgenic reporter assay in chicken embryos (Figure 7A). Six tandem copies of the *HB1RE* motif were placed upstream of a *lacZ* reporter gene to assess the transcriptional potential of this motif. We used the previously characterized autoregulatory element (*ARE*) region of the *Hoxb1* gene in this assay as a control for regulatory activity [56,64]). Electroporation of the *ARE* reporter unilaterally into one side of the neural tube shows robust reporter staining in rhombomere 4 and posterior regions (Figure 7B). Ectopic expression of *Hoxb1* in combination with the *ARE* reporter shows that HOXB1 protein can bind and enhance reporter expression, while over-expression of *Hoxb3* has no obvious effect on reporter activity. Similar analysis with the *HB1RE* reporter construct shows it is also able to direct reporter expression in the chicken neural tube, suggesting it has regulatory activity (Figure 7C). However, in contrast to the ARE vector, over-expression of *Hoxb1* in combination with the *HB1RE* reporter construct completely suppressed reporter activity. Conversely, overexpressing *Hoxb3* with the *HB1RE* reporter induced an expansion of the reporter expression domain. These results suggest that the *HB1RE* motif may be responsive to multiple HOX proteins in regulating downstream target genes in the genome. However, there appears to be a context-specific impact on activity, as the reporter data indicate HOXB1 has a repressive effect on the HB1RE motif, while HOXB3 has an activating role. These *in vivo* experiments clearly imply that the *HB1RE* motif is a novel binding site capable of receiving functional inputs from HOX proteins.

**Figure 7:**
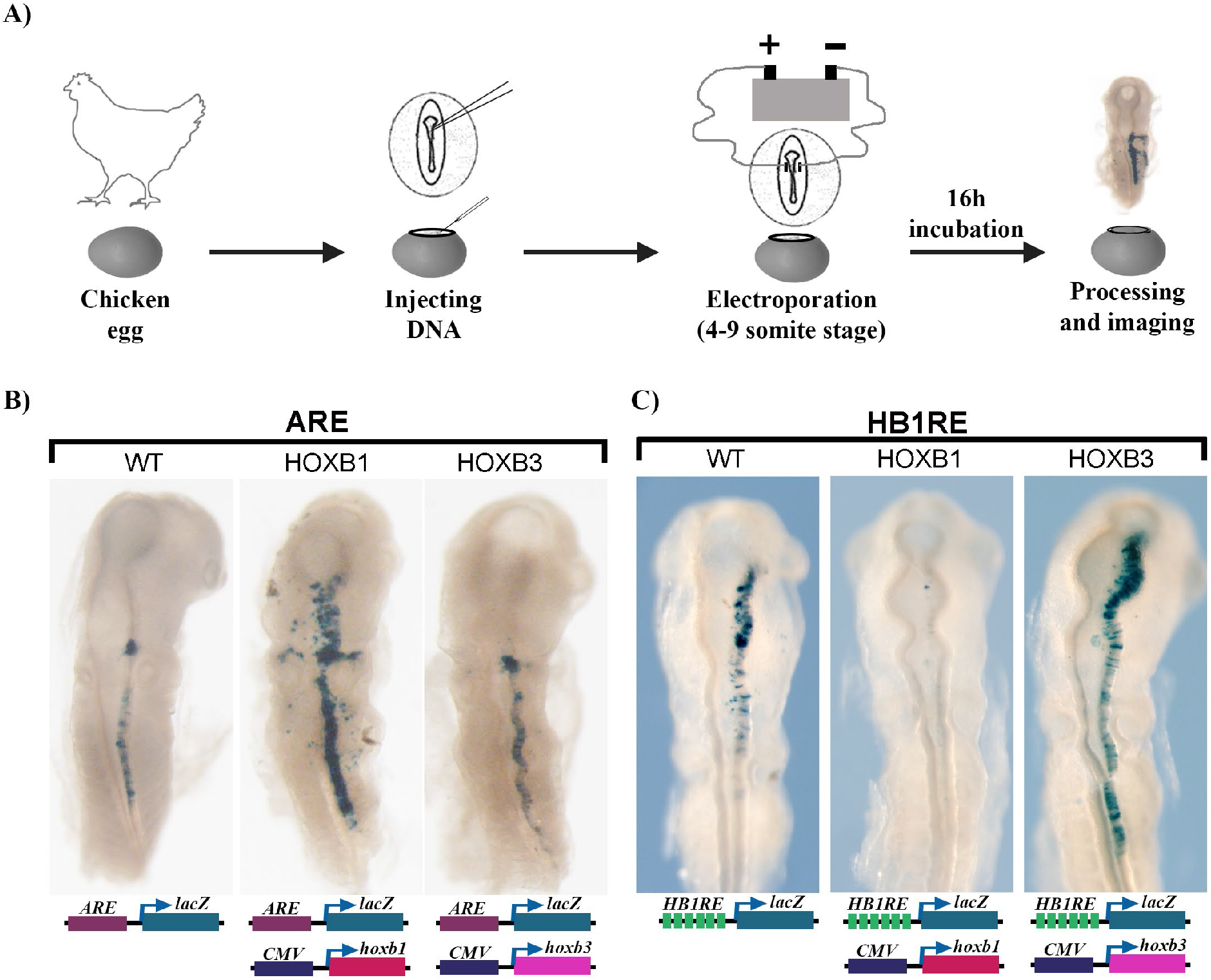
Functional analysis of HB1RE in chicken embryos: A) An outline of the experiment shows injection, electroporation of DNA in chicken embryos. B) Auto regulatory element (ARE) of HOXB1 is tested alone (left) and with CMV promoter driven overexpression of HOXB1 (middle) and HOXB3 (right). C) 6X-HB1RE motif is tested alone (left) and with CMV promoter driven overexpression of HOXB1 (middle) and HOXB3 (right).

## 4. Discussion

In this study, we have investigated the genome-wide binding properties of HOXB1 in mouse ES cells differentiated into neural fates. Using genomic, biochemical and *in vivo* regulatory assays we characterized the properties and regulatory potential a novel HOXB1 binding motif, *HB1RE*. We find that the regions bound by HOXB1 are distinctly different from those of its paralog HOXA1. The characterization of the distinct features of the HOXB1 bound peaks provides mechanistic insights into the properties associated with its functional diversification during evolution. Our findings raise several interesting questions and have implications for understanding how HOX proteins modulate the activity of downstream genes and pathways in development, disease, and evolution.

### 4.1 HOXB1 and the PBX and MEIS co-factors

Previous studies using genetic, genomic, and biochemical approaches have demonstrated the importance of interactions between HOX proteins and the co-factors PBX and MEIS, which impact their DNA binding specificities and *in vivo* functions [33,66]. These co-factor interactions appear to be highly conserved and broadly employed in a wide range of animal species [66]. Consistent with this mode for regulating the specificity of DNA binding, we have shown that HOXA1 displays significant overlaps in genome-wide co-occupancy and interactions with PBX and MEIS on its downstream target genes [26–28]. We expected that the same paradigm would generally apply to its paralog HOXB1, in light of detailed work revealing that HOXB1 partners with PBX and MEIS to directly auto- and cross-regulate segmental expression of several other *Hox* genes in the developing hindbrain [56,57,62–65]. However, the results in this study have shown that the genome-wide binding patterns of HOXB1 do not correlate well with that of PBX and MEIS. Only a small number (~7%) of the total HOXB1 binding peaks display co-occupancy with PBX and/or MEIS (Figure 1A). Furthermore, there is very little overlap between the regions bound by HOXA1 and HOXB1. The difference in binding patterns and downstream target genes is consistent with the recent report suggesting that HOXB1 has functionally diverged from HOXA1 in part through changes in a short amino motif that alters interactions with PBX proteins [26].

There are other major differences in comparing the properties of HOXB1 and HOXA1 bound regions in mouse ES cells and their differentiated derivatives. Analysis of chromatin accessibility and histone marks has shown that HOXA1 binds primarily to regions that are open based on ATAC-seq and exhibit marks characteristic of active genes [28,36]. In contrast, we find that ~ 70% of HOXB1 binding peaks are not located in accessible chromatin and lack active histone marks (Figure 1A). The small number of HOXB1 bound peaks that do display open chromatin and active marks (group 1) are co-bound by PBX and MEIS. REST occupancy defines another class (group 2) which are in open chromatin, but they contain repressive marks. The majority of HOXA1 binding peaks are located adjacent to genes [28], while we observe the majority of HOXB1 peaks positioned in gene desserts. Hence, it is not surprising that HOXA1 and HOXB1 are associated with regulating downstream target genes linked to very different cellular and developmental processes.

The difference in preference for binding to accessible chromatin domains between HOXA1 and HOXB1 is interesting in light of recent findings demonstrating that several posterior HOX proteins display differences in their abilities to bind inaccessible chromatin sites [70]. This raises the possibility that HOXB1 may bind to inaccessible regions of chromatin and in some cases partner with PBX and MEIS to enhance chromatin accessibility and gene activity. MEIS has previously been shown to have the ability to control the access of histone deacetylases to a regulatory element bound by HOX proteins [71].

The *HB1RE* motif discovered in our analysis illustrates complexities in mediating interactions between HOX proteins and PBX and MEIS. There are no obvious consensus PBX or MEIS binding sequences in the *HB1RE* motif. However, the template affinity assay demonstrated that this motif can bind PBX and MEIS proteins *in vitro* (Figure 6C). We favor the idea that the recruitment of PBX and MEIS to this template is indirect, mediated by other proteins bound to this site. Studies have shown that bipartite HOX-PBX motifs and closely positioned PBX-MEIS binding sites are effective in generating ternary complexes containing HOX, PBX and MEIS proteins on *Hox*-response elements [64,65,72]. This shows that HOX co-factors can be recruited to HOX bound regulatory regions from near adjacent sites. We also find evidence for clusters of PBX and MEIS binding sites and occupancy in HOXA1 binding peaks [27]. Hence, it is possible that indirect recruitment of PBX and MEIS to complexes with HOX proteins may be an important mechanism for bringing these proteins together to potentiate gene expression and modulate DNA binding specificity.

### 4.2 The relationship between REST, HOXB1 and repression

Transcription factors do not work alone in modulating gene expression. They frequently work in combination through the integration of synergistic, antagonist, or parallel inputs directed by the architecture and organization of DNA binding motifs in *cis*-regulatory elements. We found that a large proportion of the total HOXB1 bind peaks and ~22% of the binding peaks that contain *HB1RE* motifs display enriched motifs for REST and co-occupancy of REST proteins (Figure 1A and 3). These peaks have enrichment for a repressive histone mark and KEGG analysis shows the near adjacent genes are enriched for association with neurogenesis, neuronal processes, and behavior. The prevalence in close association of REST and HOXB1 binding motifs suggests the possibility of regulatory interactions. The REST protein complex functions as a transcriptional repressor to restrict the expression of neuronal genes in non-neural cells [67–69]. In neural development, there is evidence that REST functions to control the progressive timing of differentiation of neuronal progenitors. Hence, it would be interesting to examine whether HOXB1 impacts REST activity in neural or non-neuronal cells, by enhancing or inhibiting its inputs on shared target genes. HOXB1 can function as a positive input to stimulate gene expression on *Hox*-response elements in hindbrain segmentation [56,57,62–65]. This is illustrated by its ability to activate reporter expression from the *ARE* when electroporated into chicken embryos (Figure 7B). However, using this same assay we found that HOXB1 can also inhibit reporter expression directed by the *HB1RE*, while HOXB3 activates expression. In this context, HOXB1 displays the selective ability to repress gene expression.

### 4.3 *HB1RE* and novel enriched binding motifs

The *HB1RE* motif was one of many unknown enriched motifs detected in HOXB1 binding peaks. Our experiments provide evidence suggesting that it is a valid binding motif with regulatory potential associated with *in vivo* occupancy of HOXB1. This illustrates the challenges in trying to identify or define binding sites for TFs in the genome based on *in vitro* DNA binding specificities. There are likely to be many more non-canonical binding sites that HOX proteins and their known co-factors act upon *in vivo* and others that involve interactions with co-factors other than PBX and MEIS. While HOXA1, displayed a significant overlap with PBX and MEIS binding and enrichments for these motifs, there were many other highly enriched motifs not listed in TF DNA binding databases [27,28,36]. Hence, it is worth the effort to investigate these unknown enriched binding motifs to begin to build a better understanding of how HOX proteins are recruited to DNA and exert their regulatory activity in different developmental contexts. The emergence of improved ChIP-seq technologies that generate base pair resolution of regions bound by TFs [73], and computational pipelines to uncover and define co-associated motifs like those for REST and *HB1RE*, will help unravel the hidden grammar and context of the *cis*-regulatory code.

## Supplementary Materials

None

## Author Contributions

Conceptualization: B.D.K., N.P.S. and R.K.; genomic and proteomic experiments: B.D.K.; genomic analyses: A.P. and N.P.S.; proteomic analysis: L.F.; chicken reporter assays: M.P. and C.S.; manuscript writing, review and editing: N.P.S., B.D.K. and R.K.; All authors have read and agreed to the published version of the manuscript.

## Acknowledgments

The authors thank Ron and Joan Conaway for advice with the template affinity assay and members of the Molecular Biology, Media Preparation, Tissue Culture, Bioinformatics and Proteomics core facilities at the Stowers Institute for Medical Research for their support and help in this study. We thank other members of the Krumlauf laboratory for helpful feedback and discussion on the data and manuscript.

## References

1. Stormo, G.D.; Zhao, Y. Determining the specificity of protein-DNA interactions. Nat Rev Genet 2010, 11, 751–760, doi:10.1038/nrg2845.

2. Spitz, F.; Furlong, E.E. Transcription factors: from enhancer binding to developmental control. Nat Rev Genet 2012, 13, 613–626, doi:10.1038/nrg3207.

3. Meyer, A.; Schartl, M. Gene and genome duplications in vertebrates: the one-to-four (-to-eight in fish) rule and the evolution of novel gene functions. Curr Opin Cell Biol 1999, 11, 699–704.

4. Meyer, A.; Van de Peer, Y. From 2R to 3R: evidence for a fish-specific genome duplication (FSGD). BioEssays : news and reviews in molecular, cellular and developmental biology 2005, 27, 937–945, doi:10.1002/bies.20293.

5. Smith, J.J.; Keinath, M.C. The sea lamprey meiotic map improves resolution of ancient vertebrate genome duplications. Genome Res 2015, 25, 1081–1090, doi:10.1101/gr.184135.114.

6. Holland, P.W. Evolution of homeobox genes. Wiley interdisciplinary reviews. Developmental biology 2013, 2, 31–45, doi:10.1002/wdev.78.

7. Barrera, L.A.; Vedenko, A.; Kurland, J.V.; Rogers, J.M.; Gisselbrecht, S.S.; Rossin, E.J.; Woodard, J.; Mariani, L.; Kock, K.H.; Inukai, S., et al. Survey of variation in human transcription factors reveals prevalent DNA binding changes. Science 2016, 351, 1450–1454, doi:10.1126/science.aad2257.

8. Vaquerizas, J.M.; Kummerfeld, S.K.; Teichmann, S.A.; Luscombe, N.M. A census of human transcription factors: function, expression and evolution. Nat Rev Genet 2009, 10, 252–263, doi:10.1038/nrg2538.

9. Levine, M.; Tjian, R. Transcription regulation and animal diversity. Nature 2003, 424, 147–151.

10. Di-Poi, N.; Montoya-Burgos, J.I.; Miller, H.; Pourquie, O.; Milinkovitch, M.C.; Duboule, D. Changes in Hox genes’ structure and function during the evolution of the squamate body plan. Nature 2010, 464, 99–103, doi:10.1038/nature08789.

11. Enard, W.; Przeworski, M.; Fisher, S.E.; Lai, C.S.; Wiebe, V.; Kitano, T.; Monaco, A.P.; Paabo, S. Molecular evolution of FOXP2, a gene involved in speech and language. Nature 2002, 418, 869–872, doi:10.1038/nature01025.

12. Prud’homme, B.; Gompel, N.; Carroll, S.B. Emerging principles of regulatory evolution. Proc Natl Acad Sci U S A 2007, 104 Suppl 1, 8605–8612, doi:10.1073/pnas.0700488104.

13. Peichel, C.L.; Nereng, K.S.; Ohgi, K.A.; Cole, B.L.; Colosimo, P.F.; Buerkle, C.A.; Schluter, D.; Kingsley, D.M. The genetic architecture of divergence between threespine stickleback species. Nature 2001, 414, 901–905.

14. Noyes, M.B.; Christensen, R.G.; Wakabayashi, A.; Stormo, G.D.; Brodsky, M.H.; Wolfe, S.A. Analysis of homeodomain specificities allows the family-wide prediction of preferred recognition sites. Cell 2008, 133, 1277–1289, doi:10.1016/j.cell.2008.05.023.

15. Carroll, S.B. Evolution at two levels: on genes and form. PLoS Biol 2005, 3, e245.

16. Berger, M.F.; Badis, G.; Gehrke, A.R.; Talukder, S.; Philippakis, A.A.; Pena-Castillo, L.; Alleyne, T.M.; Mnaimneh, S.; Botvinnik, O.B.; Chan, E.T., et al. Variation in homeodomain DNA binding revealed by high-resolution analysis of sequence preferences. Cell 2008, 133, 1266–1276, doi:10.1016/j.cell.2008.05.024.

17. Nitta, K.R.; Jolma, A.; Yin, Y.; Morgunova, E.; Kivioja, T.; Akhtar, J.; Hens, K.; Toivonen, J.; Deplancke, B.; Furlong, E.E., et al. Conservation of transcription factor binding specificities across 600 million years of bilateria evolution. eLife 2015, 4, doi:10.7554/eLife.04837.

18. Jolma, A.; Yan, J.; Whitington, T.; Toivonen, J.; Nitta, K.R.; Rastas, P.; Morgunova, E.; Enge, M.; Taipale, M.; Wei, G., et al. DNA-binding specificities of human transcription factors. Cell 2013, 152, 327–339, doi:10.1016/j.cell.2012.12.009.

19. Pearson, J.C.; Lemons, D.; McGinnis, W. Modulating Hox gene functions during animal body patterning. Nat Rev Genet 2005, 6, 893–904, doi:10.1038/nrg1726.

20. Carroll, S.B. Homeotic genes and the evolution of arthropods and chordates. Nature 1995, 376, 479–485, doi:10.1038/376479a0.

21. Carroll, S.B.; Weatherbee, S.D.; Langeland, J.A. Homeotic genes and the regulation and evolution of insect wing number. Nature 1995, 375, 58–61, doi:10.1038/375058a0.

22. Deschamps, J.; Duboule, D. Embryonic timing, axial stem cells, chromatin dynamics, and the Hox clock. Genes Dev 2017, 31, 1406–1416, doi:10.1101/gad.303123.117.

23. Tschopp, P.; Duboule, D. A genetic approach to the transcriptional regulation of Hox gene clusters. Annu Rev Genet 2011, 45, 145–166, doi:10.1146/annurev-genet-102209-163429.

24. Parker, H.J.; Krumlauf, R. Segmental arithmetic: summing up the *Hox* gene regulatory network for hindbrain development in chordates. Wiley interdisciplinary reviews. Developmental biology 2017, 6, doi:10.1002/wdev.286.

25. Mallo, M.; Wellik, D.M.; Deschamps, J. Hox genes and regional patterning of the vertebrate body plan. Dev Biol 2010, 344, 7–15, doi:10.1016/j.ydbio.2010.04.024.

26. Singh, N.P.; De Kumar, B.; Paulson, A.; Parrish, M.E.; Zhang, Y.; Florens, L.; Conaway, J.W.; Si, K.; Krumlauf, R. A six-amino-acid motif is a major determinant in functional evolution of HOX1 proteins. Genes Dev 2020, 34, 1680–1696, doi:10.1101/gad.342329.120.

27. De Kumar, B.; Parker, H.J.; Paulson, A.; Parrish, M.E.; Pushel, I.; Singh, N.P.; Zhang, Y.; Slaughter, B.D.; Unruh, J.R.; Florens, L., et al. HOXA1 and TALE proteins display cross-regulatory interactions and form a combinatorial binding code on HOXA1 targets. Genome Res 2017, 27, 1501–1512, doi:10.1101/gr.219386.116.

28. De Kumar, B.; Parker, H.J.; Paulson, A.; Parrish, M.E.; Zeitlinger, J.; Krumlauf, R. Hoxa1 targets signaling pathways during neural differentiation of ES cells and mouse embryogenesis. Dev Biol 2017, 432, 151–164, doi:10.1016/j.ydbio.2017.09.033.

29. Moens, C.B.; Selleri, L. Hox cofactors in vertebrate development. Dev Biol 2006, 291, 193–206.

30. Mann, R.S.; Lelli, K.M.; Joshi, R. Hox specificity unique roles for cofactors and collaborators. Curr Top Dev Biol 2009, 88, 63–101, doi:10.1016/S0070-2153(09)88003-4.

31. Dard, A.; Reboulet, J.; Jia, Y.; Bleicher, F.; Duffraisse, M.; Vanaker, J.M.; Forcet, C.; Merabet, S. Human HOX Proteins Use Diverse and Context-Dependent Motifs to Interact with TALE Class Cofactors. Cell reports 2018, 22, 3058–3071, doi:10.1016/j.celrep.2018.02.070.

32. Slattery, M.; Riley, T.; Liu, P.; Abe, N.; Gomez-Alcala, P.; Dror, I.; Zhou, T.; Rohs, R.; Honig, B.; Bussemaker, H.J., et al. Cofactor binding evokes latent differences in DNA binding specificity between Hox proteins. Cell 2011, 147, 1270–1282, doi:10.1016/j.cell.2011.10.053.

33. Merabet, S.; Mann, R.S. To Be Specific or Not: The Critical Relationship Between Hox And TALE Proteins. Trends in genetics : TIG 2016, 32, 334–347, doi:10.1016/j.tig.2016.03.004.

34. Mann, R.; Chan, S.-K. Extra specificity from extradenticle: the partnership between HOX and PBX/EXD homeodomain proteins. Trends in Genetics 1996, 12, 258–262.

35. Beard, C.; Hochedlinger, K.; Plath, K.; Wutz, A.; Jaenisch, R. Efficient method to generate single-copy transgenic mice by site-specific integration in embryonic stem cells. Genesis 2006, 44, 23–28, doi:10.1002/gene.20180.

36. De Kumar, B.; Parker, H.J.; Parrish, M.E.; Lange, J.J.; Slaughter, B.D.; Unruh, J.R.; Paulson, A.; Krumlauf, R. Dynamic regulation of Nanog and stem cell-signaling pathways by Hoxa1 during early neuro-ectodermal differentiation of ES cells. Proc Natl Acad Sci U S A 2017, 114, 5838–5845, doi:10.1073/pnas.1610612114.

37. Sheikh, B.N.; Downer, N.L.; Kueh, A.J.; Thomas, T.; Voss, A.K. Excessive versus Physiologically Relevant Levels of Retinoic Acid in Embryonic Stem Cell Differentiation. Stem Cells 2014, 32, 1451–1458, doi:10.1002/stem.1604.

38. Smith, K.T.; Martin-Brown, S.A.; Florens, L.; Washburn, M.P.; Workman, J.L. Deacetylase inhibitors dissociate the histone-targeting ING2 subunit from the Sin3 complex. Chemistry & biology 2010, 17, 65–74, doi:10.1016/j.chembiol.2009.12.010.

39. De Kumar, B.; Parrish, M.E.; Slaughter, B.D.; Unruh, J.R.; Gogol, M.; Seidel, C.; Paulson, A.; Li, H.; Gaudenz, K.; Peak, A., et al. Analysis of dynamic changes in retinoid-induced transcription and epigenetic profiles of murine Hox clusters in ES cells. Genome Res 2015, 25, 1229–1243, doi:10.1101/gr.184978.114.

40. Buenrostro, J.D.; Wu, B.; Chang, H.Y.; Greenleaf, W.J. ATAC-seq: A Method for Assaying Chromatin Accessibility Genome-Wide. Current protocols in molecular biology / edited by Frederick M. Ausubel ... [et al.] 2015, 109, 21 29 21–29, doi:10.1002/0471142727.mb2129s109.

41. Langmead, B.; Salzberg, S.L. Fast gapped-read alignment with Bowtie 2. Nature methods 2012, 9, 357–359, doi:10.1038/nmeth.1923.

42. Zhang, Y.; Liu, T.; Meyer, C.A.; Eeckhoute, J.; Johnson, D.S.; Bernstein, B.E.; Nusbaum, C.; Myers, R.M.; Brown, M.; Li, W., et al. Model-based analysis of ChIP-Seq (MACS). Genome Biol 2008, 9, R137, doi:10.1186/gb-2008-9-9-r137.

43. Gupta, S.; Stamatoyannopoulos, J.A.; Bailey, T.L.; Noble, W.S. Quantifying similarity between motifs. Genome Biol 2007, 8, R24, doi:10.1186/gb-2007-8-2-r24.

44. Kel-Margoulis, O.V.; Kel, A.E.; Reuter, I.; Deineko, I.V.; Wingender, E. TRANSCompel: a database on composite regulatory elements in eukaryotic genes. Nucleic Acids Res 2002, 30, 332–334, doi:10.1093/nar/30.1.332.

45. Khan, A.; Fornes, O.; Stigliani, A.; Gheorghe, M.; Castro-Mondragon, J.A.; van der Lee, R.; Bessy, A.; Cheneby, J.; Kulkarni, S.R.; Tan, G., et al. JASPAR 2018: update of the open-access database of transcription factor binding profiles and its web framework. Nucleic Acids Res 2018, 46, D1284, doi:10.1093/nar/gkx1188.

46. Bailey, T.L.; Elkan, C. Fitting a mixture model by expectation maximization to discover motifs in biopolymers. Proceedings / ... International Conference on Intelligent Systems for Molecular Biology; ISMB. International Conference on Intelligent Systems for Molecular Biology 1994, 2, 28–36.

47. Ogata, H.; Goto, S.; Sato, K.; Fujibuchi, W.; Bono, H.; Kanehisa, M. KEGG: Kyoto Encyclopedia of Genes and Genomes. Nucleic Acids Res 1999, 27, 29–34, doi:10.1093/nar/27.1.29.

48. Gottschalk, A.J.; Timinszky, G.; Kong, S.E.; Jin, J.; Cai, Y.; Swanson, S.K.; Washburn, M.P.; Florens, L.; Ladurner, A.G.; Conaway, J.W., et al. Poly(ADP-ribosyl)ation directs recruitment and activation of an ATP-dependent chromatin remodeler. Proc Natl Acad Sci U S A 2009, 106, 13770–13774, doi:10.1073/pnas.0906920106.

49. Sela, D.; Chen, L.; Martin-Brown, S.; Washburn, M.P.; Florens, L.; Conaway, J.W.; Conaway, R.C. Endoplasmic reticulum stress-responsive transcription factor ATF6alpha directs recruitment of the Mediator of RNA polymerase II transcription and multiple histone acetyltransferase complexes. The Journal of biological chemistry 2012, 287, 23035–23045, doi:10.1074/jbc.M112.369504.

50. Florens, L.; Washburn, M.P. Proteomic analysis by multidimensional protein identification technology. Methods Mol Biol 2006, 328, 159–175, doi:10.1385/1-59745-026-X:159.

51. Washburn, M.P.; Wolters, D.; Yates, J.R., 3rd. Large-scale analysis of the yeast proteome by multidimensional protein identification technology. Nat Biotechnol 2001, 19, 242–247, doi:10.1038/85686.

52. Tabb, D.L.; McDonald, W.H.; Yates, J.R., 3rd. DTASelect and Contrast: tools for assembling and comparing protein identifications from shotgun proteomics. J Proteome Res 2002, 1, 21–26.

53. Gould, A.; Itasaki, N.; Krumlauf, R. Initiation of rhombomeric *Hoxb4* expression requires induction by somites and a retinoid pathway. Neuron. 1998, 21, 39–51.

54. Inoue, T.; Krumlauf, R. An impulse to the brain: Using in vivo electroporation. Nature Neuroscience 2001, 4, 6–8.

55. Itasaki, N.; Bel-Vialar, S.; Krumlauf, R. ‘Shocking’ developments in chick embryology: electroporation and in ovo gene expression. Nature cell biology 1999, 1, E203–207, doi:10.1038/70231.

56. Popperl, H.; Bienz, M.; Studer, M.; Chan, S.K.; Aparicio, S.; Brenner, S.; Mann, R.S.; Krumlauf, R. Segmental expression of Hoxb-1 is controlled by a highly conserved autoregulatory loop dependent upon exd/pbx. Cell 1995, 81, 1031–1042.

57. Maconochie, M.K.; Nonchev, S.; Studer, M.; Chan, S.K.; Popperl, H.; Sham, M.H.; Mann, R.S.; Krumlauf, R. Cross-regulation in the mouse *HoxB* complex: the expression of *Hoxb2* in rhombomere 4 is regulated by *Hoxb1*. Genes and Development 1997, 11, 1885–1896.

58. Studer, M.; Lumsden, A.; Ariza-McNaughton, L.; Bradley, A.; Krumlauf, R. Altered segmental identity and abnormal migration of motor neurons in mice lacking *Hoxb-1*. Nature 1996, 384, 630–634, doi:10.1038/384630a0.

59. Arenkiel, B.R.; Tvrdik, P.; Gaufo, G.O.; Capecchi, M.R. Hoxb1 functions in both motoneurons and in tissues of the periphery to establish and maintain the proper neuronal circuitry. Genes Dev 2004, 18, 1539–1552.

60. Webb, B.D.; Shaaban, S.; Gaspar, H.; Cunha, L.F.; Schubert, C.R.; Hao, K.; Robson, C.D.; Chan, W.M.; Andrews, C.; MacKinnon, S., et al. HOXB1 founder mutation in humans recapitulates the phenotype of Hoxb1 -/- mice. Am J Hum Genet 2012, 91, 171–179, doi:10.1016/j.ajhg.2012.05.018.

61. McClintock, J.M.; Kheirbek, M.A.; Prince, V.E. Knockdown of duplicated zebrafish hoxb1 genes reveals distinct roles in hindbrain patterning and a novel mechanism of duplicate gene retention. Development 2002, 129, 2339–2354.

62. Tümpel, S.; Cambronero, F.; Ferretti, E.; Blasi, F.; Wiedemann, L.M.; Krumlauf, R. Expression of *Hoxa2* in rhombomere 4 is regulated by a conserved cross-regulatory mechanism dependent upon *Hoxb1*. Developmental Biology 2007, 302, 646–660.

63. Studer, M.; Gavalas, A.; Marshall, H.; Ariza-McNaughton, L.; Rijli, F.M.; Chambon, P.; Krumlauf, R. Genetic interactions between Hoxa1 and Hoxb1 reveal new roles in regulation of early hindbrain patterning. Development 1998, 125, 1025–1036.

64. Ferretti, E.; Cambronero, F.; Tümpel, S.; Longobardi, E.; Wiedemann, L.M.; Blasi, F.; Krumlauf, R. *Hoxb1* enhancer and control of rhombomere 4 expression: Complex interplay between PREP1-PBX1-HOXB1 binding sites. Molecular and cellular biology 2005, 25, 8541–8552.

65. Ferretti, E.; Marshall, H.; Pöpperl, H.; Maconochie, M.; Krumlauf, R.; Blasi, F. Segmental expression of Hoxb2 in r4 requires two separate sites that integrate cooperative interactions between Prep1, Pbx and Hox proteins. Development 2000, 127, 155–166.

66. Hudry, B.; Thomas-Chollier, M.; Volovik, Y.; Duffraisse, M.; Dard, A.; Frank, D.; Technau, U.; Merabet, S. Molecular insights into the origin of the Hox-TALE patterning system. eLife 2014, 3, e01939, doi:10.7554/eLife.01939.

67. Mandel, G.; Fiondella, C.G.; Covey, M.V.; Lu, D.D.; Loturco, J.J.; Ballas, N. Repressor element 1 silencing transcription factor (REST) controls radial migration and temporal neuronal specification during neocortical development. Proc Natl Acad Sci U S A 2011, 108, 16789–16794, doi:10.1073/pnas.1113486108.

68. Ballas, N.; Mandel, G. The many faces of REST oversee epigenetic programming of neuronal genes. Curr Opin Neurobiol 2005, 15, 500–506, doi:10.1016/j.conb.2005.08.015.

69. Ballas, N.; Grunseich, C.; Lu, D.D.; Speh, J.C.; Mandel, G. REST and its corepressors mediate plasticity of neuronal gene chromatin throughout neurogenesis. Cell 2005, 121, 645–657, doi:10.1016/j.cell.2005.03.013.

70. Bulajic, M.; Srivastava, D.; Dasen, J.S.; Wichterle, H.; Mahony, S.; Mazzoni, E.O. Differential abilities to engage inaccessible chromatin diversify vertebrate Hox binding patterns. Development 2020, 147, doi:10.1242/dev.194761.

71. Choe, S.K.; Lu, P.; Nakamura, M.; Lee, J.; Sagerstrom, C.G. Meis cofactors control HDAC and CBP accessibility at Hox-regulated promoters during zebrafish embryogenesis. Dev Cell 2009, 17, 561–567, doi:10.1016/j.devcel.2009.08.007.

72. Jacobs, Y.; Schnabel, C.A.; Cleary, M.L. Trimeric association of Hox and TALE homeodomain proteins mediates *Hoxb2* hindbrain enhancer activity. Molecular and cellular biology 1999, 19, 5134–5142.

73. He, Q.; Johnston, J.; Zeitlinger, J. ChIP-nexus enables improved detection of in vivo transcription factor binding footprints. Nat Biotechnol 2015, 33, 395–401, doi:10.1038/nbt.3121.

